## Supplementary figures and images for "Open the LID: LXRα regulates ChREBPα transactivity in a target gene-specific manner through an agonist-modulated LBD-LID interaction"

### Supplementary figures

Suppl. Figure 1

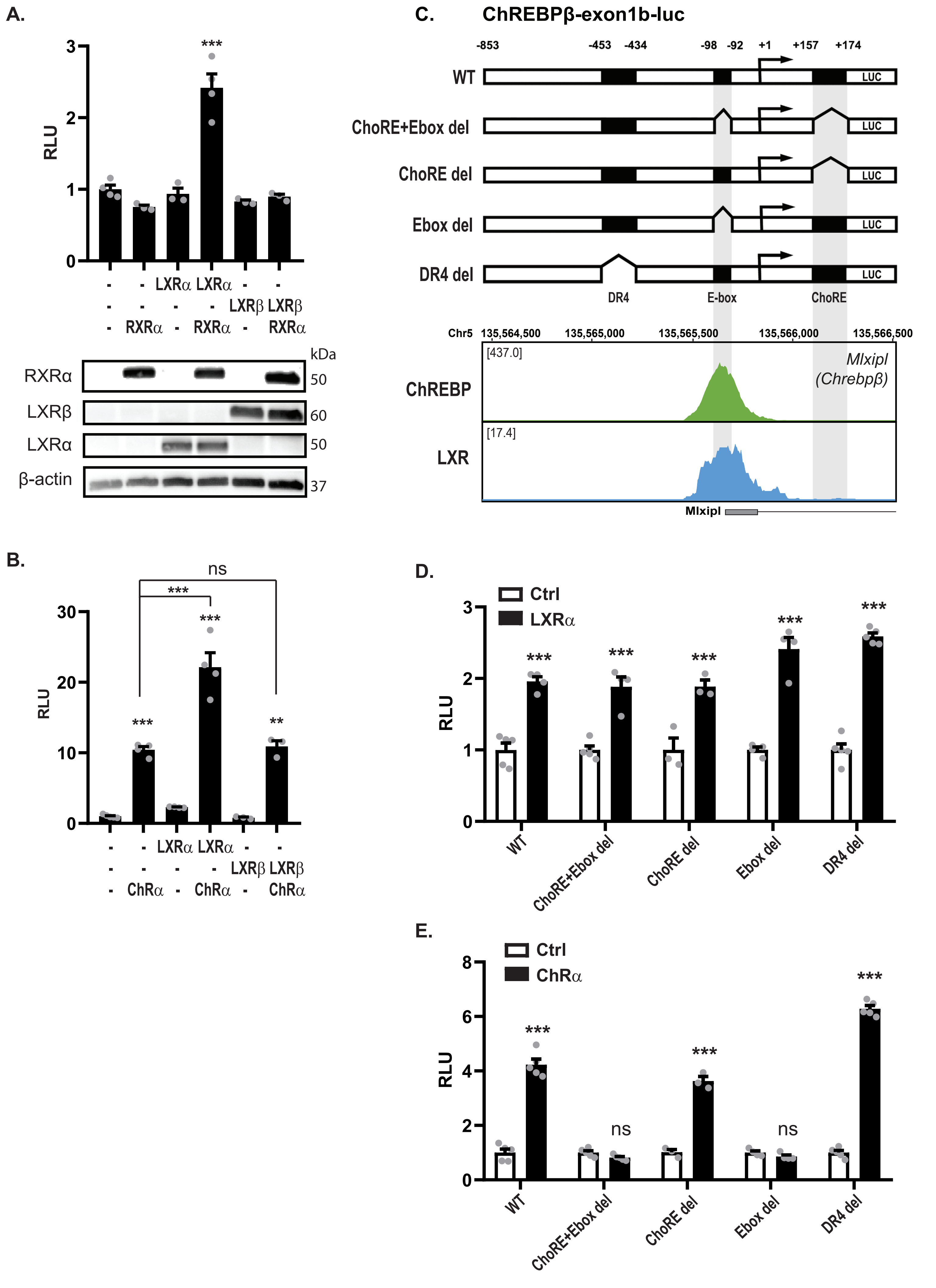

##### Suppl. Figure 2

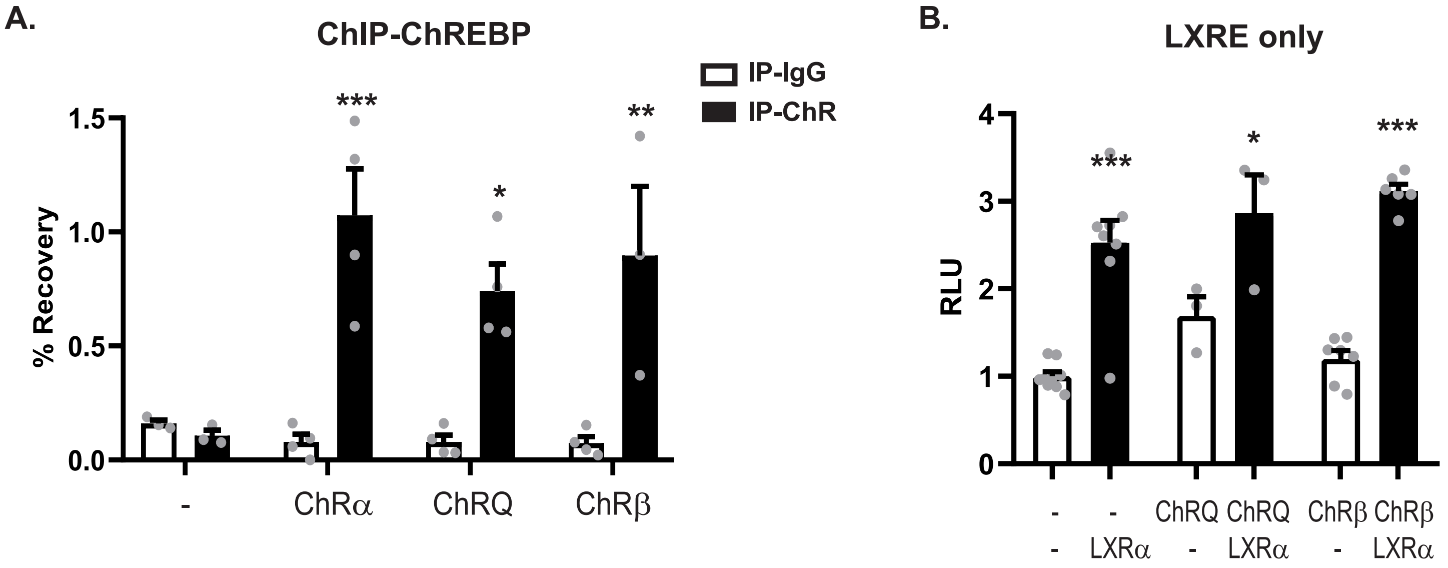


##### Suppl. Figure 3

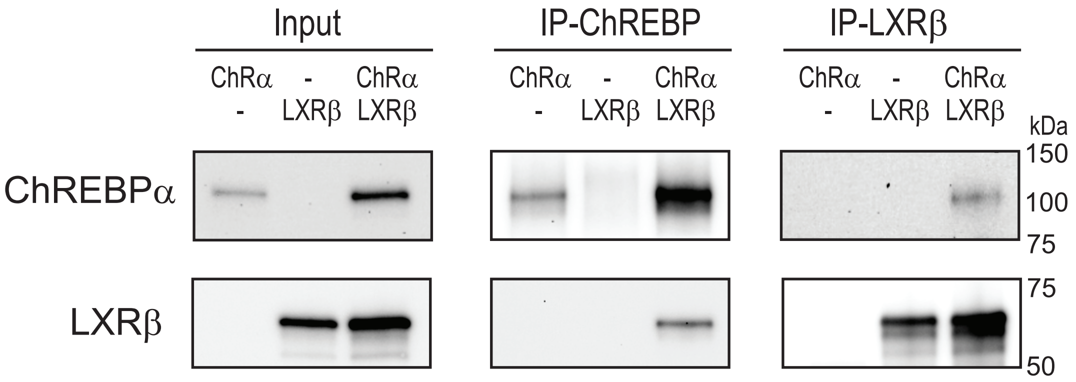


##### Suppl. Figure 4

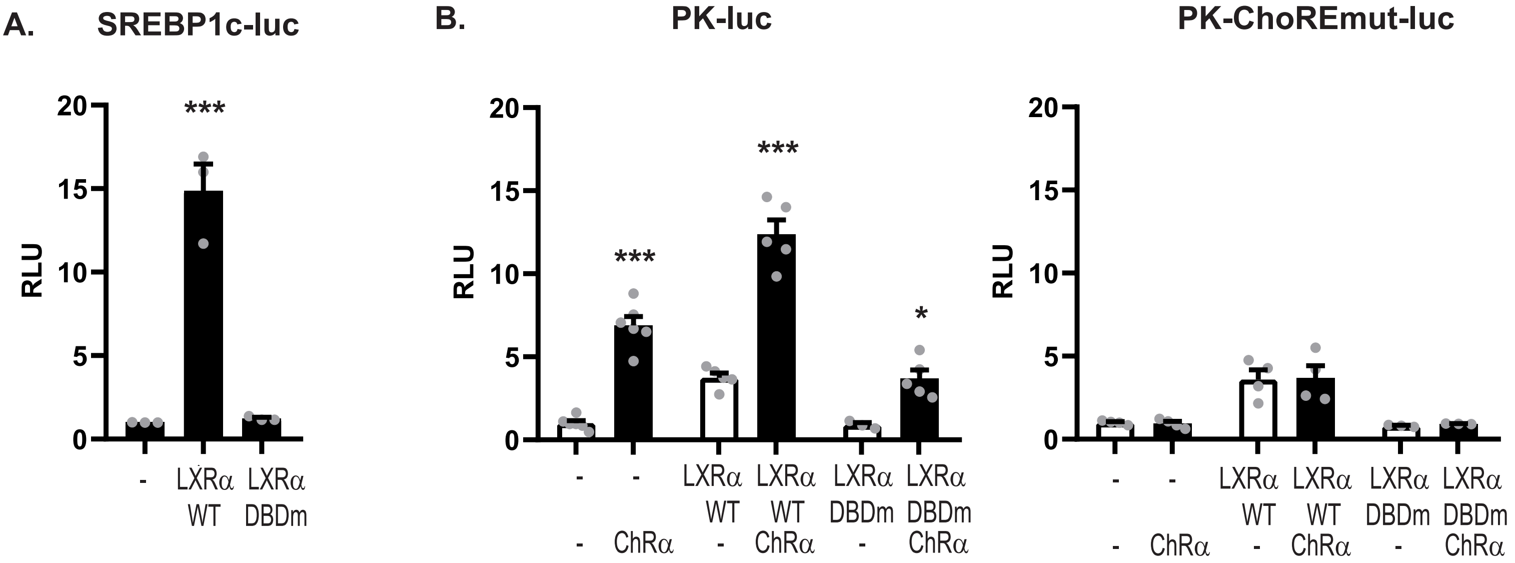

#### Suppl. Figure 5

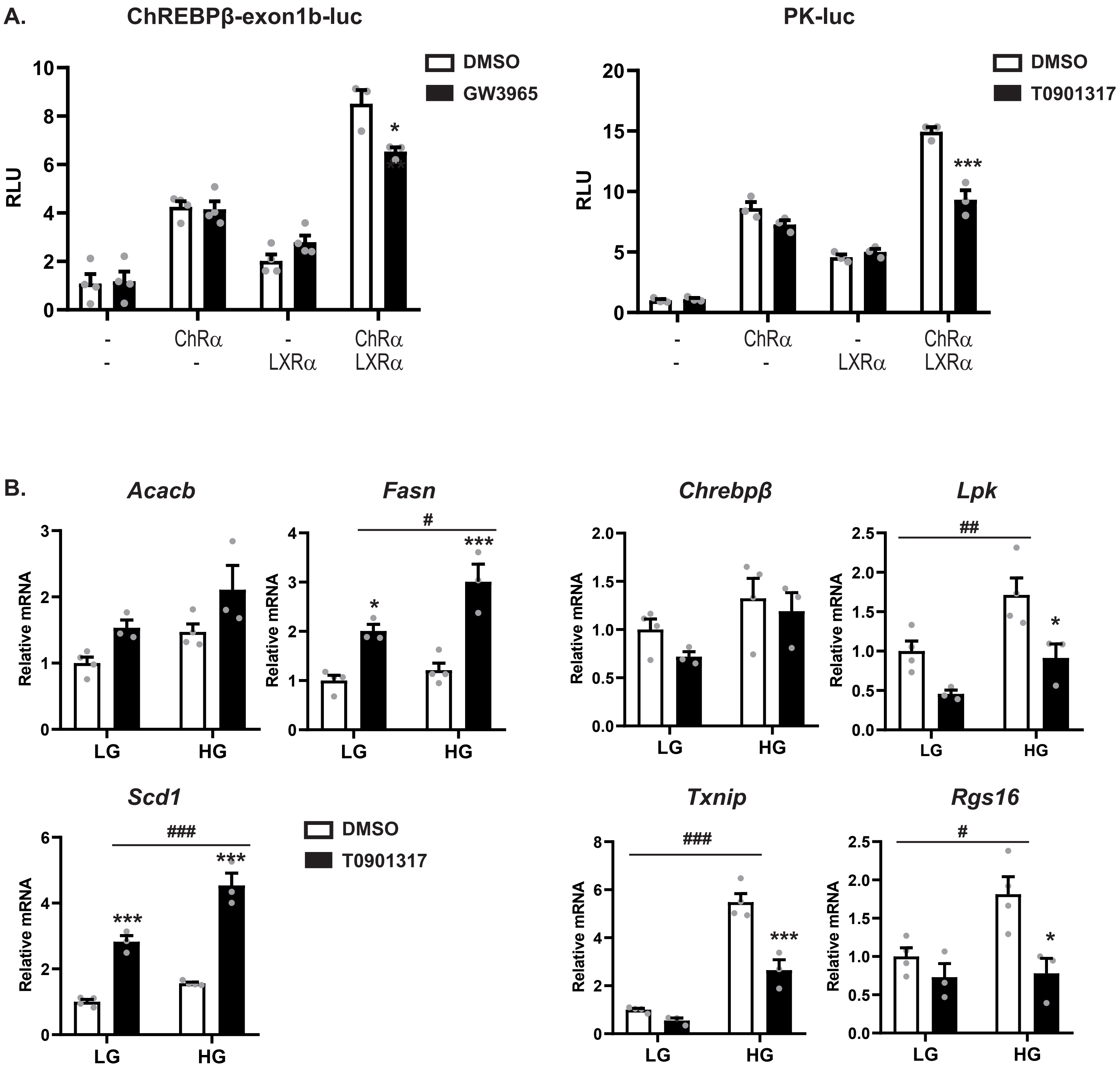

Suppl. Figure 6

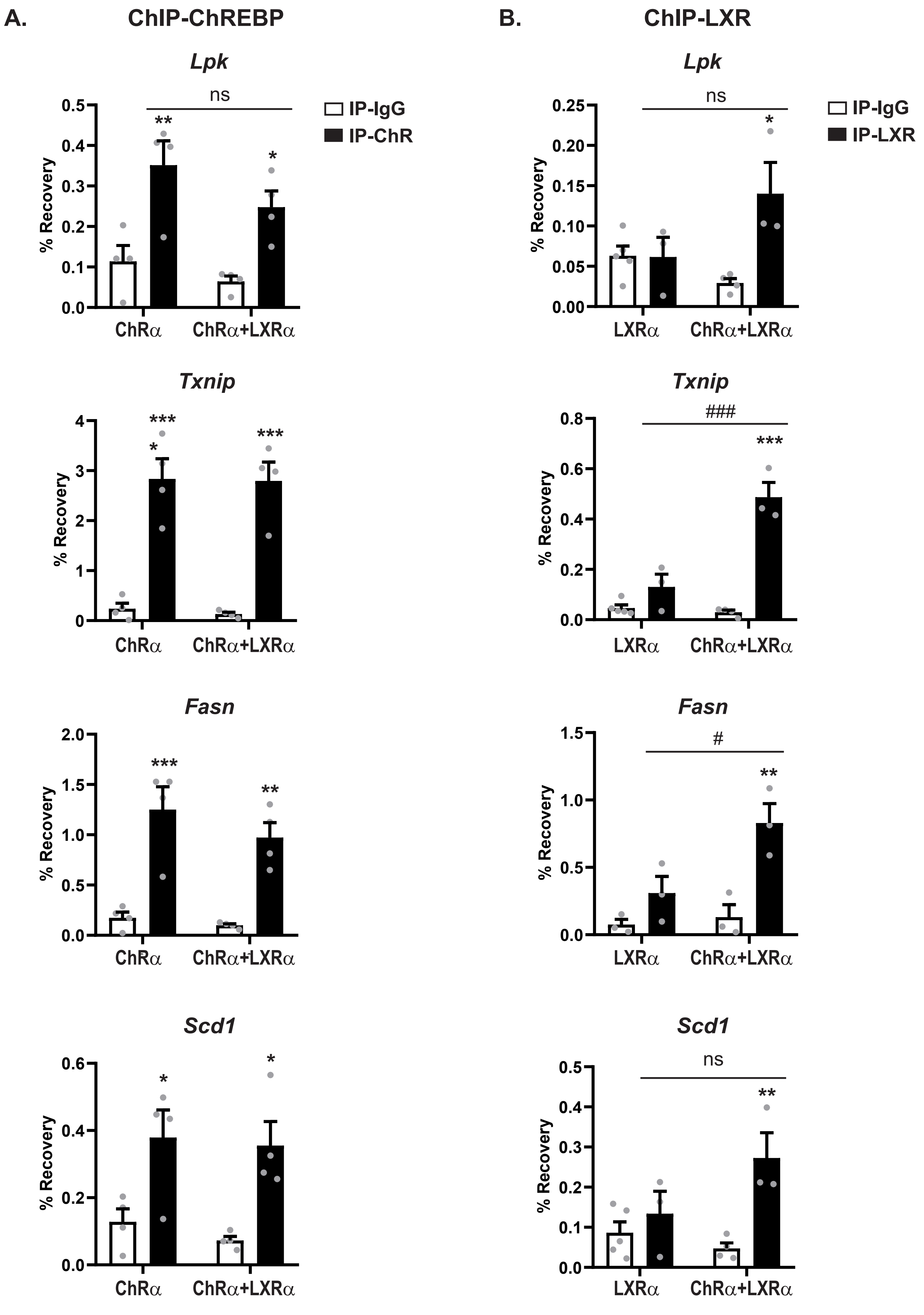
